# Genome-wide analysis of Corsican population reveals a close affinity with Northern and Central Italy

**DOI:** 10.1101/722165

**Authors:** Erika Tamm, Julie Di Cristofaro, Stéphane Mazières, Erwan Pennarun, Alena Kushniarevich, Alessandro Raveane, Ornella Semino, Jacques Chiaroni, Luisa Pereira, Mait Metspalu, Francesco Montinaro

## Abstract

Despite being the fourth largest island in the Mediterranean basin, the genetic variation of Corsica has not been explored as exhaustively as Sardinia, which is situated only 11 km South. However, it is likely that the populations of the two islands shared, at least in part, similar demographic histories. Moreover, the relative small size of the Corsica may have caused genetic isolation, which, in turn, might be relevant under medical and translational perspectives. Here we analysed genome wide data of 16 Corsicans, and integrated with newly (33 individuals) and previously generated samples from West Eurasia and North Africa. Allele frequency, haplotype-based, and ancient genome analyses suggest that although Sardinia and Corsica may have witnessed similar isolation and migration events, the latter is genetically closer to populations from continental Europe, such as Northern and Central Italians.

## Introduction

Corsica, located south of the shore of Côte d’Azur (France), and west of Tuscany (Italy), is separated from Sardinia to its south by the Strait of Bonifacio. It is the fourth largest Mediterranean island and unlike most of them, its relief is very mountainous, with a mountain range bisecting the island.

The understanding of the peopling of Corsica has remained incomprehensive. From a geological perspective, during the last glaciation, Corsica and Sardinia formed a single landmass and its distance to Italy was reduced, possibly increasing connections with mainland^1,2^. Furthermore, archaeological records suggest that the Southern part of the Sardinia-Corsica palaeo-island, characterised by milder climate and less geographical asperities, was settled at a first stage, with the area corresponding to modern day Corsica, characterised by harsher conditions, being colonised later. However, the acidity of deposits and submersion led to a scarce persistence of anthropological and archaeological remains, preventing the extensive characterization of its peopling dynamics. There is no clear evidence of human traces from the end of Pleistocene and the beginning of Holocene. The oldest human remains found so far in Corsica are from Campu Stefano and are dated 8,940 ^14^C year BP (10216–9920 cal. BP, 95.4% range)^3^. Archaeological and genetic data suggest episodic and discontinuous settlements during Mesolithic and transitional phases between Mesolithic and Neolithic periods^4,5^. The permanent human presence in Corsica is attested in the Neolithic period since the 6^th^ millennium BC. From this period interactions with mainland and other islands are illustrated with wider appearance of non-local lithic resources and development of similar ceramic traditions over a larger Western Mediterranean region^6^.

In the last three millennia, the Corsican population witnessed several dramatic reductions of population size due to conquests, epidemics outbreaks and economic crises. The Greeks established the city of Alalia (today Aleria) in 565 BC, which is also the first mention of Corsica in historical records. Subsequently, it witnessed numerous intrusions and conquests by different populations. Carthaginians and Etruscans dominated the island until the Roman occupation in the third century BC. Successive invasions by the Vandals, Ostrogoths and Saracens, took place until the beginning of the Byzantine influence in the 6^th^ century, followed by Roman Papacy from the 8^th^ century onwards. From the end of the 11^th^ century Corsica was under the government of Pisa. Then, with the intermittent period dominated by the kings of Aragon, Corsica was under the rule of Genoa from the 13^th^ to the 18^th^ century. After a short period of independence, Corsica was incorporated into France in 1768 through the Treaty of Versailles^7^.

Previous genetic studies on Corsican population have shown regional differentiation^8–11^, such as a marked North-South differentiation, confirmed by surname and linguistic studies^12^, and mirroring the geographic features of the island.

Despite their geographic proximity, different surveys based on unilinear and autosomal markers have produced contradicting results in characterizing the relationship between Sardinia and Corsica. Some studies have found genetic affinity between the two islands^8,9,13–15^, while conversely, some have suggested genetic distinction^10,11,16,17^. Furthermore, the extent of genetic relationship with mainland populations is controversial. Close genetic connections have been observed between Corsica and continental Mediterranean populations^10,17–19^, whereas other studies have reported limited genetic influences from mainland^9,11^. Some studies have pointed out that different regions in Corsica have received genetic influences from distinct sources, showing diverse affinities with surrounding populations^8,11^.

Regardless of its genetic affinity, it is unclear to what extent the Corsican population shows a signature of demographic bottleneck or population decrease, as expected in a long-term isolation scenario. Previous studies have demonstrated that isolated populations are valuable resources in genome-wide association studies. In geographically and/or culturally isolated conditions evolutionary forces and population dynamics can lead to the high level of homozygosity, reduced genetic diversity and increase in peculiar allele frequencies, which makes easier to trace genetic variants affecting medical or phenotypic traits^20^. Sardinia is a well-known example (see for instance^21^ and references therein). Preliminary studies have suggested Corsican potential in these investigations: an analysis on chromosome X microsatellite markers demonstrated an overall LD decrease in the innermost part of Corsica^22^ and some medical and association studies have been conducted on Corsican population^23–25^. However, it was pointed out that caution should be taken in making general assumptions^22^, underlining the importance of characterising the population under investigation before designing and planning of gene association studies^26–28^.

Despite its anthropological and epidemiological relevance, a genome-wide characterisation of Corsican population is not available so far. The aims of the current study were to explore the internal genetic structure of Corsicans using the genome-wide data, and to assess the genetic relation to neighbouring modern and ancient European populations. In doing so, we have genotyped 16 Corsican samples collected from different locations of the island. To better contextualize our results, we have additionally genotyped 33 new samples from Portugal, France and Italy and combined this newly generated data with 892 modern and 222 ancient Eurasian and African genomes from previously published sources (Supplementary Table S1).

## Results

### Population Structure

To explore whether Corsicans, as insular population, display characteristic traces of isolation and endogamy in their genomes, we assessed Runs of Homozygosity (RoH). RoHs are uninterrupted segments of homozygous genotypes present in individuals. Their number and extension are correlated to the level of identity by descent and are largely affected by population’s demographic history^29,30^. Corsicans show an excess of RoHs indicated by the high median values of total number and total length of RoHs, next to Sardinians and French Basques (Fig. 1). This homozygosity pattern is characteristic of isolated populations with relatively low effective population size (Ne) and high degree of endogamy^29–31^. Similar to Sardinians and Basques, Corsicans show high variability of long RoHs, possibly indicating recent on-going admixture.

**Figure 1.**
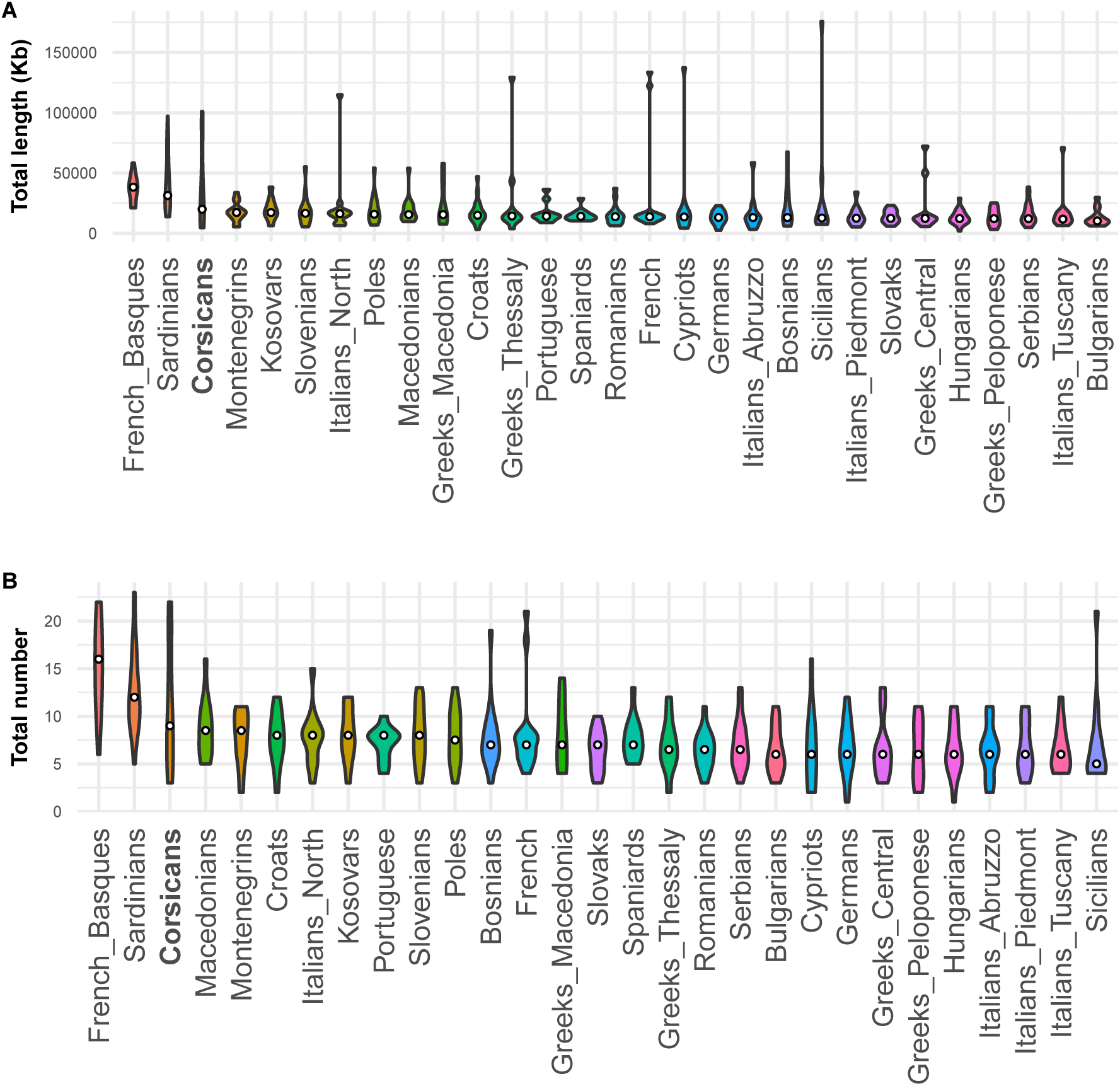
Violin plots of Runs of Homozygosity (RoH) estimations for Corsicans and comparative European populations. A) Total length in kilobases (kb) of genome in RoH. B) Total number of RoH segments. The violin plots show the distribution of RoH fragments with width indicating frequencies and the median as a white circle. Populations are ordered from left to right according to their median value.

To position Corsicans in the context of their geographic neighbours we used an unsupervised clustering approach^32^ implemented in the program ADMIXTURE^33^. At the level of lowest cross-validation index (K=6), Corsicans and geographically related populations are composed mainly by three ancestry components -“*Sardinian*”, *“Northern and Eastern Europe”*, “*Caucasus and Middle East”* (dark blue, light blue, and lime green respectively; Fig. 2, Supplementary Fig. S1, S2). Corsicans are most similar to North-Central Italian populations (Piedmont, Lombardy, Tuscany), displaying a slightly larger proportion of a modal component in Sardinians. This similarity in ADMIXTURE profiles remains throughout higher levels of K values, even when further components appear (Fig. 2, Supplementary Fig. S1). The closest geographic neighbours, Sardinians, and the other known isolate, French Basques, virtually lack Caucasus/Middle Eastern (lime green) ancestry. At K = 8 and K=11 two components almost fixed in Sardinians (dark blue) and in French Basques (middle blue), respectively, emerge.

**Figure 2.**
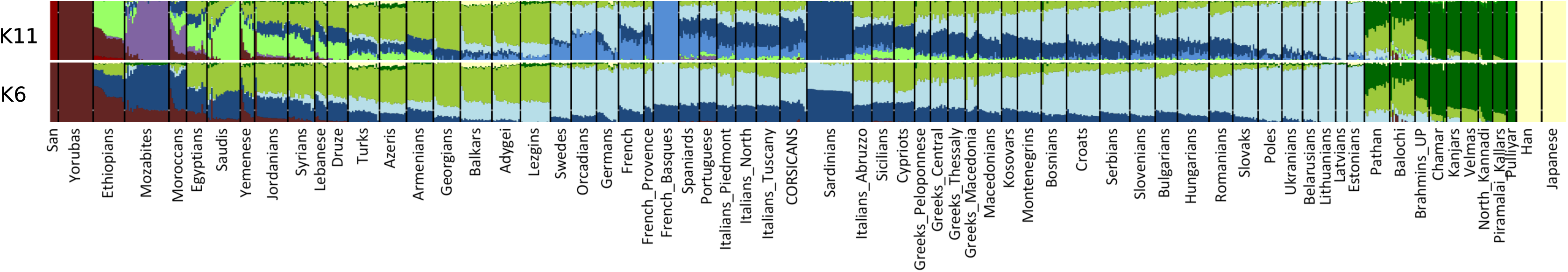
ADMIXTURE plot of individual ancestry proportions at K = 6 and K = 11.

Analysis of average population pairwise F_st_ distance confirms ADMIXTURE results showing shortest genetic distances between geographically close mainland populations and revealing larger distances with Sardinians and French Basques (Supplementary Table S2).

To evaluate the genetic affinity between Corsica and neighbouring populations we carried out f3-statistics analysis. Outgroup f3 statistics measures the amount of shared genetic drift between two populations from an outgroup^34^. The results showed high affinity of Corsicans with Sardinians and French Basques, followed, as with the ADMIXTURE and F_ST_ analyses, by Northern Italians and a series of mainland European populations (Supplementary Fig. S3A). The tests of admixture based on the f3 statistics revealed an overall complexity in the admixture history of Corsica. Out of 2,652 tests performed, 18 were statistically significant (Supplementary Fig. S3B), and most of them described Corsica as a combination of Sardinian and Caucasus/Northern European contributions. Moreover, six tests including populations from North African/Arabia and one from Northern Europe were statistically significant.

### Haplotype-based analyses

We inferred the fine-scale population structure harnessing the haplotype-sharing patterns among individuals. In detail, using ChromoPainter we reconstructed each analysed individual *j* as a mosaic of genomic fragments inherited by *n* donor samples^35^. The resulting *i x n* coancestry chunkcount matrix was then employed to identify and characterise the homogenous groups of individuals, in the form of a dendrogram. fineSTRUCTURE clustering results together with the ChromoPainter coancestry chunkcount matrix were subsequently used to identify, date and describe admixture events with GLOBETROTTER^36^.

The complete fineSTRUCTURE dendrogram is represented in Supplementary Fig. S4, together with detailed chunkcount coancestry matrix (Supplementary Fig. S5) and pairwise coincidence matrix (Supplementary Fig. S6). For simplicity, we assigned a name to the clusters summarising their composition, reported in Supplementary Table S3. All together, we identified 85 homogeneous groups, with a broad correlation with geographic origin of samples. Specifically, West Eurasian subjects grouped into 38 clusters, for which six macrogroups may be identified (Fig. 3A). Samples from the Levant area (Druze, Syrians_Lebanese and Lebanese clusters) grouped close to individuals from Cyprus and Armenia in the Caucasus (purple in Fig. 3). Other samples from Caucasus (light blue in Fig. 3) fell into a macrogroup that includes eight different clusters (Lezgins, Azeris, Turks, Georgians, Balkars_Adygei, Balkars, Adygei1, Adygei2). Samples from North and West Europe distributed into six clusters (FrenchBasques, French, Iberia2, Orcadians1, Orcadians2, Swedes), which grouped together and were nearby a group of seven clusters comprising mostly individuals from the Balkans (CentralEastEurope, Balkan1, Kosovars, Montenegrins, Balkan2, Greeks, Bosnian_Greek). These two macrogroups were closest to the Central and East European group, which was composed by four clusters (Baltics, Germans, Estonians, EasternEurope). Finally, the only clusters solely represented by Corsicans (Supplementary Fig. 4) grouped with all the Italian samples and one Iberian cluster (Iberia1), establishing a Southern European macrogroup.

**Figure 3.**
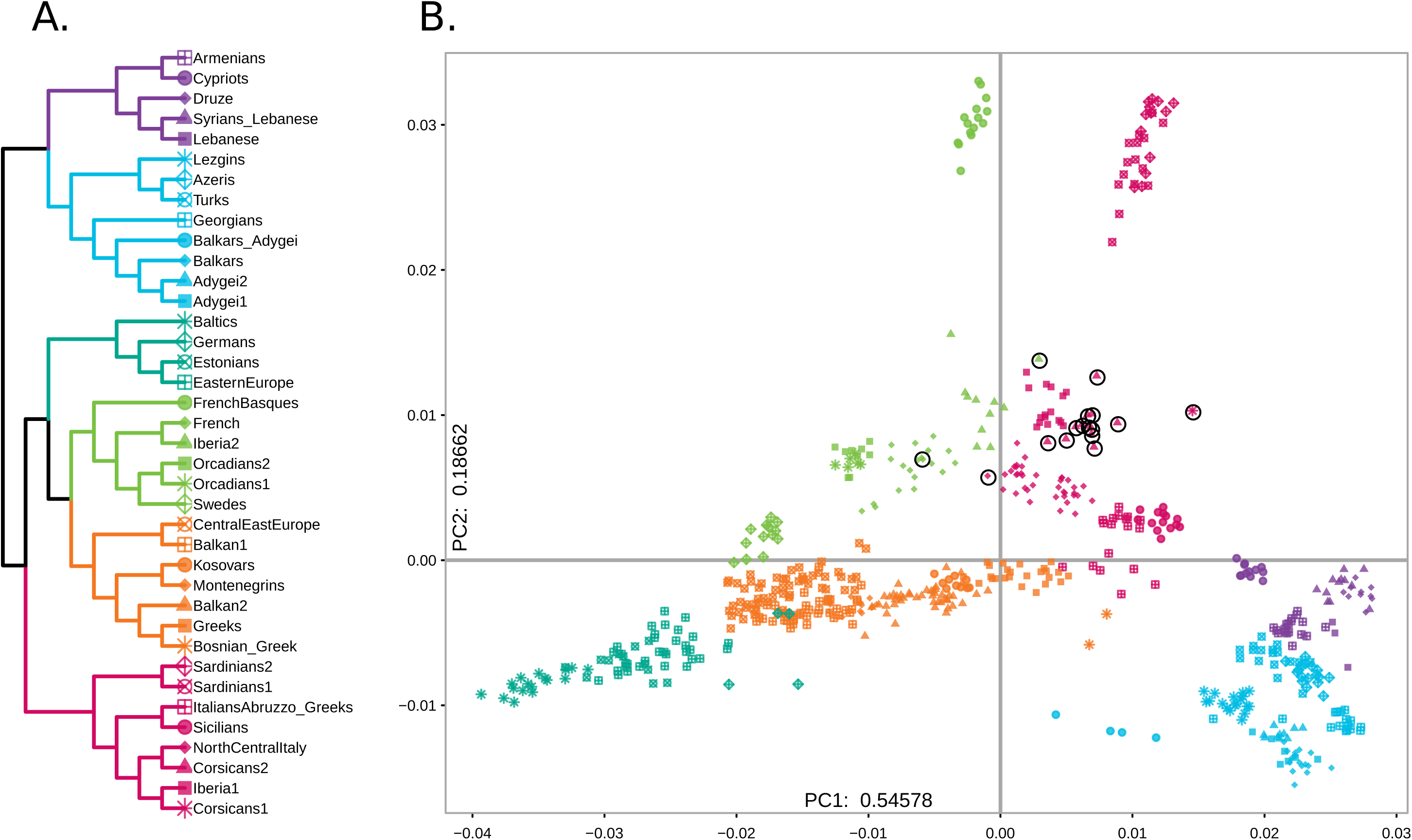
**A) fineSTRUCTURE tree of Eurasian populations.** We applied the ChromoPainter/fineStucture pipeline in order to build a dendrogram showing the relationships among homogenous groups in our dataset. Only the portion of the tree including Western Eurasian and a sub-set of Levantine populations are shown. Colours of the tree branches indicate macroregional grouping discussed in the text. The full fineSTRUCTURE tree is shown in Supplementary Figure S4. **B) Principal Component Analysis based on haplotype sharing.** The chunkcount coancestry matrix of Western Eurasian and Levantine samples have been used to perform PCA analysis, and visualize the top two components in a scatterplot. Corsican samples are highlighted by black circles.

**Figure 4.**
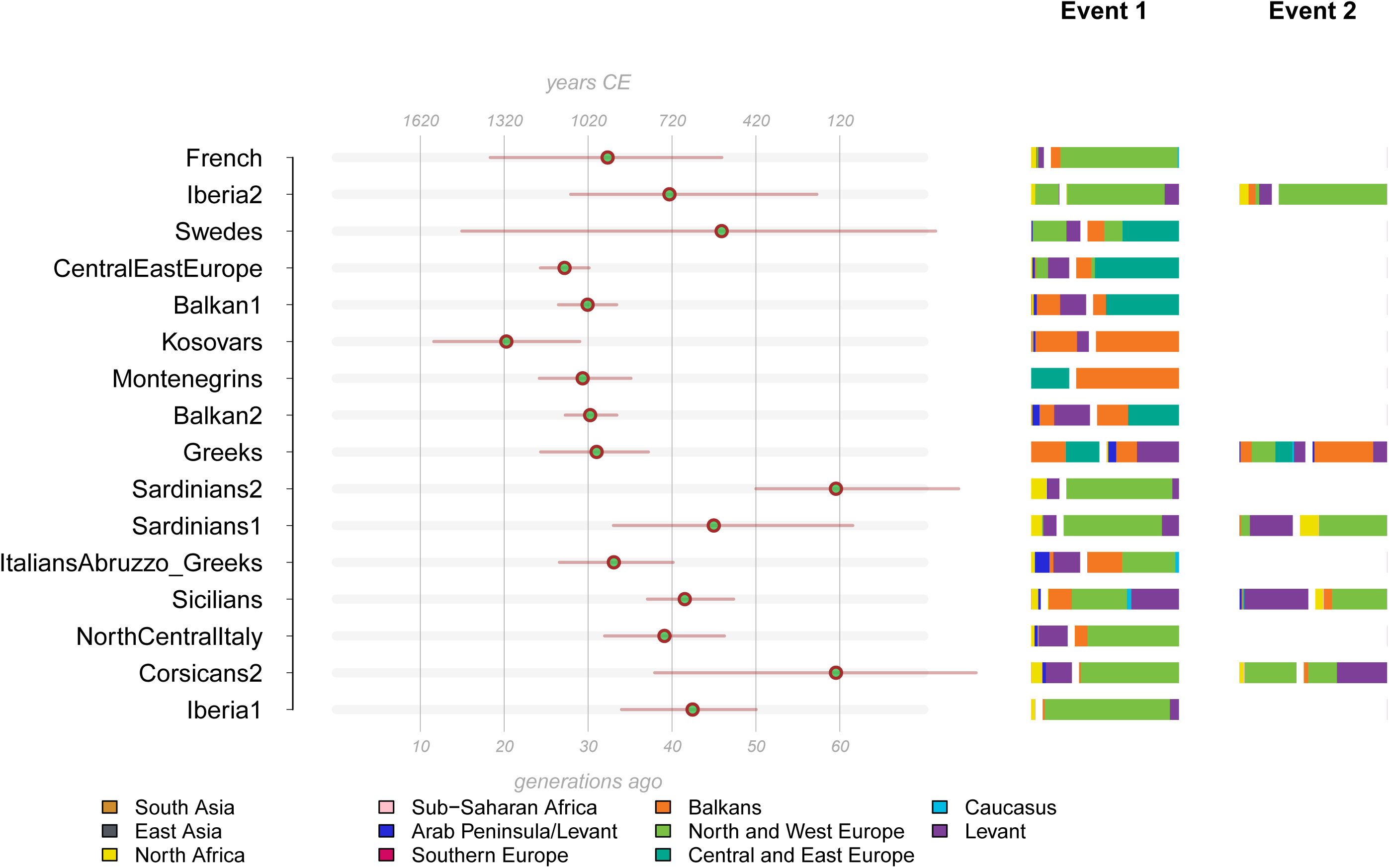
Admixture dates as inferred by GLOBETROTTER in the “non-local” analysis. We fit the painting profile of Western Eurasian populations into expected curves for different admixture models, as implemented in GLOBETROTTER. The estimated dates and sources composition are shown. When one date multiple way result was detected, two events are indicated. For each event the two putative source composition is separated by a white space in the barplot.

At a finer inspection, out of the 16 Corsican samples analysed, 12 formed a population specific cluster (Corsicans2) in the Southern European clade, related to North and Central Italian group. Three samples fell into three different clusters composed by North and Central Italians (NorthCentralItaly), French (French) and Iberians (Iberia2), respectively (Fig. 3, Supplementary Fig. S4), possibly reflecting recent relationship with mainland populations. The remaining Corsican individual formed a separate branch, closest to Portuguese and Spaniards (Iberia1). Sardinians and French Basques group with neighbouring populations: the former with Southern Europeans while the latter with Northern and Western Europeans.

In order to explore the relationships between populations, we performed a Principal Component Analysis (PCA) based on the chunkcount coancestry matrix. The first two PCs explain a large proportion of the total variance (55% and 19% respectively, Fig. 3B), confirming the efficacy in summarizing the genomic information of the painting approach. The first principal component separates populations along a North-South axis, placing North-East Europeans on one side (left) and Near East/Caucasus populations to the opposite (right). The second component separates populations along a West-East axis, with Sardinians and French Basques being clear outliers. Most Corsican samples group together close to Italian and Spanish populations; the four samples that did not group with Corsican main cluster, occupy outlier positions also in the PCA.

To estimate the admixture history of Corsican population, genetic clusters defined by fineSTRUCTURE were used to perform analysis with GLOBETROTTER^36^. We focused on Western Eurasian clusters composed by more than five individuals. As geographically close populations tend to share recent ancestry and distant genetic contacts may be masked, we performed two different GLOBETROTTER analyses, “full” and “non-local”, as previously reported^37,38^ (Fig. 4, Supplementary Fig. S7 and Table S4). The “full” analysis considers all samples as possible sources, while “non-local” excludes Southern European clusters as donors. In both the analysis, a single admixture involving more than one source was identified for the main Corsican cluster. This admixture involved North and West Europe and Levant/North African sources, and occurred between 37 and 76 generations ago, a time period spanning the fall of the Roman Empire and the invasions by Barbarians and Saracens.

In the “full” analysis, the admixture sources included Sardinians, NorthCentral Italians, Spanish and Sicilians (Supplementary Table S4), possibly suggesting gene-flows from continental Europe, while the Levant/North African contribution inferred in the “non-local” analysis was most probably passed to Corsicans hitchhiking on mainland populations. Similar admixture profiles were observed for two Sardinian clusters, with central time estimates of 59 and 44 generations ago. These results suggest that similar admixture episodes affected both the Mediterranean islands. The impact of North Africa and Levant is also evident in the remaining Italian and Iberian samples, highlighting the wide impact of the event.

### Ancient contribution in Corsica

In order to understand how Corsica and neighbour populations are related to groups that occupied the continent in the last ∼10k years, we have performed a Principal Component Analysis projecting ancient individuals onto PCs estimated on modern European allele-frequency variation (Fig. 5A). When the first two PCs are considered, Corsicans are close to European samples from Early, Middle Neolithic and Chalcolithic, together with Balkans Chalcolithic and Neolithic. Compared to other relevant modern European populations, Corsicans show their closeness to Central and Northern Italian rather than Sardinians, although they are scattered towards the latter, suggesting a larger affinity to Neolithic ancestry and Sardinian population than mainland European.

**Figure 5.**
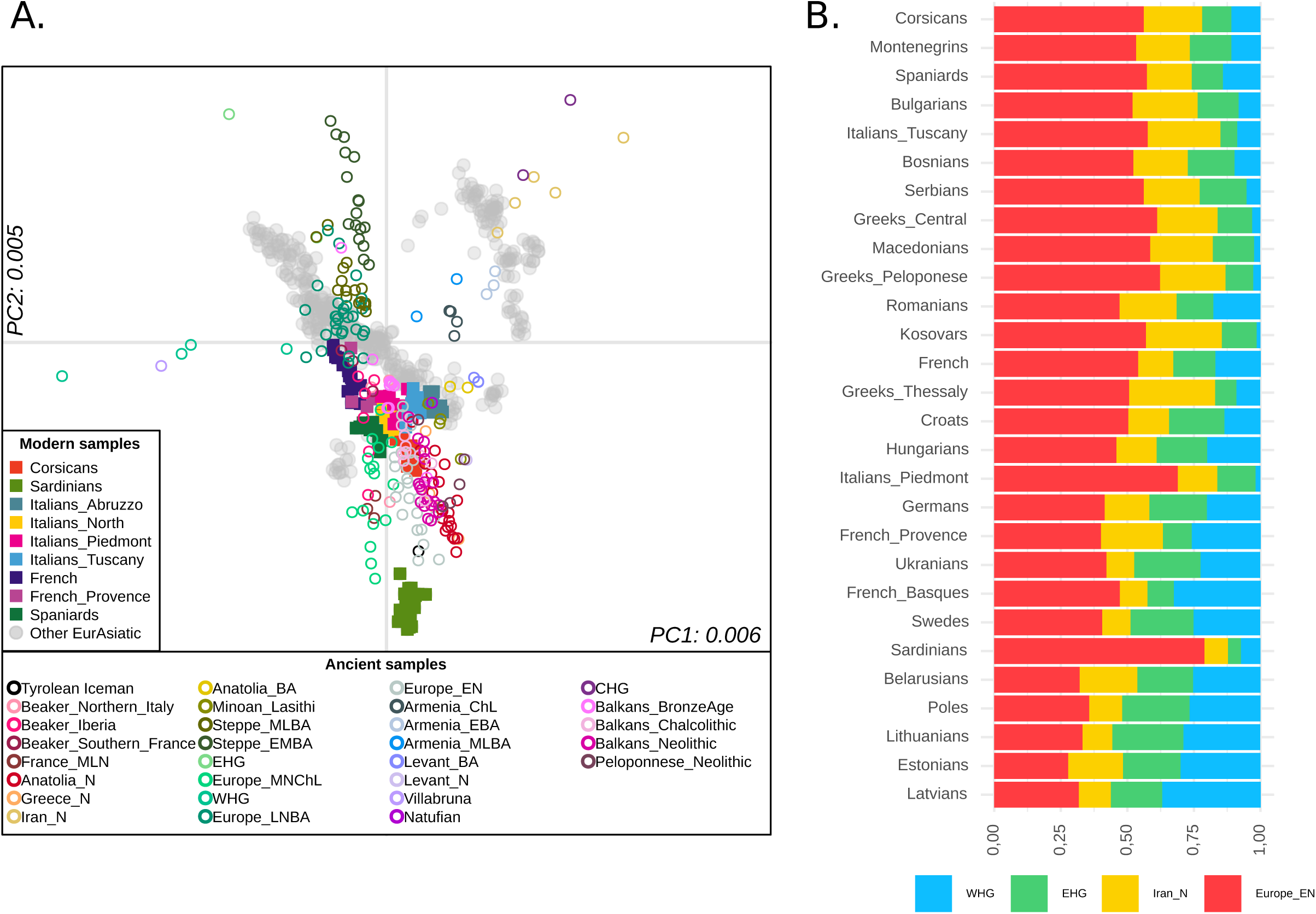
A) Principal Component Analysis of ancient individual genotypes projected onto the first two PCs estimated using modern West Eurasian populations. **B) Admixture profile of Western Eurasian as inferred by qpAdm.** We have reconstructed the admixture profile of all the analysed populations with qpADM, which harnesses a combination of f4 describing the relationship of “target” and “sources” with a set of outgroups. A four-population model including WHG, EHG, European Early Neolithic (EN) and Iran Neolithic is supported in most of the tested populations. Populations are sorted according to Euclidean distances.

In order to better characterize the “ancestral” composition of Corsican and European populations, we inferred their genetic relationship with a set of ancient individuals using qpAdm^39^. In our analysis, the Corsican samples were characterised by a high ancestry of European Early Neolithic (56%), similarly to Italian, Spanish and Balkan populations (Fig. 5B). In addition, Corsica harboured a relatively high proportion related to Iranian Neolithic (22%), while the contribution of Western and Eastern Hunter Gatherer (WHG, EHG) was smaller, about 11%. According to previous researches, a substantial proportion of the EHG and Iranian Neolithic (related to Caucasus Hunter Gatherer, CHG) trace back to Bronze Age movements from the Steppe, however, Iranian Neolithic could have arrived to Western Mediterranean with different migration^38–40^. This seems to be supported also for Corsica when qpAdm analysis including Steppe_EMBA (Early Middle Bronze Age) and Iranian Neolithic (Iran_N) were considered in the same analysis evaluating the proportions (Anatolia Neolithic: 33%, Steppe_EMBA: 19%, Iran Neolithic: 14%, Europe Middle Neolithic/Chalcolithic: 34%). Compared to Corsicans, French and Spanish samples were characterised by a smaller proportion of Iranian Neolithic (13% and 15%, respectively), and a slightly higher contribution from Western Hunter Gatherers (14% and 17%).

## Discussion

Despite being the fourth largest island in the Mediterranean basin, the genetic variation of Corsica has not been explored as exhaustively as Sardinia^41^, which is situated only 11 km South. However, it is likely that the two populations have shared at least part of their demographic history, given their geographic proximity and similarities in the archaeological records. In addition, the relatively small size of Corsica may have contributed to create isolation conditions affecting the genetic variability of the autochthonous population^22^.

Our analysis revealed that Corsican population shares several genomic features with Sardinia and North-Central Italy, creating a unique blend of genomic ancestries (Figs. 1-3).

The Corsican population shows a relatively high homozygosity characterised by high variance, suggesting that it witnessed both an isolation and migration phase. A similar pattern has been observed for Sardinians (Fig. 1). This result confirms the importance of Corsican population as a valuable resource for association studies.

Allele frequency-based assignment algorithm and genetic distance methods showed that Corsica is genetically more related to mainland populations from France, Italy, Spain and Greece rather than Sardinia (Fig. 2, Supplementary Fig. S1, Supplementary Table S2). In contrast, the f3 outgroup analysis suggested that Corsica shares a high amount of genetic drift with Sardinia and Basque, followed by North Italy (Fig. S3A).

In order to better characterise the genetic relationship of Corsica with other Mediterranean populations we have harnessed the information embedded in haplotype configurations. The analyses of the haplotype sharing patterns performed using ChromoPainter and fineSTRUCTURE provided substantial evidences of higher affinity between Corsica and Central or Northern Italian populations rather than Sardinian, French and Iberian populations (Fig. 3). In fact, most of the Corsican individuals tested fall in a homogenous cluster closely related to North and Central Italy. On the other hand, a small proportion of individuals are closer to French and Iberians, suggesting the existence of substantial heterogeneity possibly due to recent migration, as confirmed by RoHs analysis (Fig. 1).

When we investigated admixture evidence of Corsican population through f3 analysis, Corsicans could be described as a combination of allele frequencies from Sardinia, Northern Europe, Caucasus, North Africa and Arabian Peninsula (Fig. S3B). A similar result was obtained when haplotypic patterns were explored (Fig. 4, Supplementary Fig. S7 and Table S4). Corsicans fit a scenario of admixture involving more than two sources related to Southern European, Western European and Levant Arabic populations which occurred ∼60 generations ago, in a similar time frame inferred for Sardinians, although the large confidence intervals associated with the analysis make the overall interpretation challenging. Nevertheless, similar source compositions have been inferred not only for Sardinians, but also for North and Central Italians, suggesting that similar processes may have impacted the Mediterranean basin. In detail, we inferred that admixture events involving Southern European samples occurred about between 37 and 76 generations ago, a time period spanning the fall of the Roman Empire and the invasions from Barbarians and Saracens (Fig. 4). These estimates are in accordance with those inferred in previous investigations for a sub-sample of circum-Mediterranean populations^36,37^.

Lastly, we have evaluated the genetic relationship of Corsica and other Mediterranean populations with ancient Eurasian individuals through PCA and qpAdm (Fig. 5). Corsicans show a high affinity to European ancient individuals characterized by high Neolithic ancestry such as European from Early, Middle and Late (Chalcolithic) Neolithic. Furthermore, most of the European tested populations can be modelled according to European Early Neolithic, Iranian Neolithic, Western and Eastern Hunter Gatherer contribution. The ancient profile of Corsica is characterised by a high proportion of ancestry related to European Early Neolithic, although lower than the one found in Sardinians (56% vs 79%), and similar to the one inferred for Tuscany and Spain. Interestingly, Corsica has a non-negligible fraction of ancestry related to Iranian Neolithic, which could be independent from the one brought to Europe through the Steppe related migration, as previously suggested^38,42^. In fact, we have found a similar proportion of Steppe Bronze Age (∼19%) and Iranian Neolithic (∼14%) in Corsicans.

In conclusion, the genetic characterisation of Corsica is consistent with a closer genetic affinity with Northern and Central Italian populations rather than Sardinians, although sharing with the latter a noteworthy proportion of ancestry and similar demographic and isolation processes. The analysis of larger sample sizes from different regions of the island and genetic material from ancient specimen may help to further evaluate the existence of genetic structure in the island and its demographic history.

## Materials and Methods

### Sampling

A total of 49 DNA samples were genotyped for the current study. Sixteen Corsican samples from different locations were selected for genotyping from a larger dataset based on DNA quality criteria and have been analysed previously in Y-chromosome surveys^11,43,44^. To extend the comparative dataset, additional samples were involved: ten Portuguese, five French samples from Provence, eleven Italian samples from Piedmont and seven Italian samples from Tuscany. Samples from Piedmont and Tuscany have been studied for Y chromosome^45^ and Tuscany samples also for mitochondrial DNA^46^. DNA samples have been collected from healthy unrelated individuals and all donors have provided informed consent. Experiments were carried out in accordance with the relevant guidelines and regulations of collaborative institutions. The research has been approved by the Research Ethics Committee of the University of Tartu.

### Genome-wide SNP data

DNA was extracted from blood/saliva samples and genotyping was carried out on Illumina 660K platform (Human660W-Quad BeadChip). New samples were combined with data from previous studies^51,57–64^. In total, 892 individuals from 67 populations were analysed (Supplementary Table S1). The merged dataset was preprocessed with PLINK v1.9^47^ in order to include only autosomal SNPs with minor allele frequency >0.005% and genotyping success >97%. The cryptic relationships between samples (relatives of 1st and 2nd degree) were controlled with software KING v1.4^48^ and two samples (one Yemen and one North Kannadi) were randomly removed from detected relative pairs. For some analyses SNPs in strong linkage disequilibrium (pairwise genotypic correlation R^2^ >0.4) in a window of 1,000 SNPs, sliding the window by step of 25 SNPs, were excluded. Exact numbers of individuals, populations and markers used in each analysis are specified in Supplementary Table S1.

### Runs of homozygosity

Runs of homozygosity (RoH) were inferred using PLINK v1.9^47^, with sliding window of 50 SNPs (5,000 kb), allowing for one heterozygous and five missing calls per window. RoH were defined as regions of at least 50 consecutive homozygous SNPs spanning at least 1,500 kb, with a gap of less than 1,000 kb between adjacent regions. The required minimum density was set at 50 kb/SNP^30,49^.

### ADMIXTURE

Maximum likelihood unsupervised clustering algorithm implemented in ADMIXTURE^33^ was used to infer putative ancestral components in the Corsican population in a worldwide context. Clustering was performed 100 times at K=2 to K=15 (Supplementary Fig. S1). Convergence between runs was assessed using log-likelihood scores (LL). According to a low level of variation in LL scores (LLs < 1) within the top 10% fraction of runs with the highest LLs, the global maximum was assumed to be reached at K=2 to K=6, K=9, K=11 and K=13 (Supplementary Fig. S2B). The lowest cross-validation (CV) index, which points to the predictive accuracy of the model at a given K, was observed at K=6 (Supplementary Fig. S2A).

### F_ST_

Mean population pairwise F_ST_ were calculated according to method of Weir and Cockerham 1984^50^ using a R script as in^51^.

### f3 tests

f3 tests were performed with ADMIXTOOLS v. 4.1^34^ on a subset of West Eurasian and North African populations. To remove outlier samples from included populations, a PC analysis was performed, and samples not clustering with their population on the PC plot were excluded (Supplementary Table S1). The outgroup f3 test of the form f3(Corsicans, X; Yoruba) was implemented to measure the amount of shared drift between Corsicans and other populations, with the Yoruba population from Nigeria being set as the outgroup. To test for evidence of admixture in the Corsican population as target, standard f3 statistics were computed using the test configuration f3(Corsicans; X, Y). Negative values of f3 statistics with z-score below-3 indicate statistically significant admixture in a target population.

### ChromoPainter and fineSTRUCTURE

The haplotype-based structure of Corsicans and other European populations was explored by applying the ChromoPainter/fineSTRUCTURE pipeline^35^. First, genotype data was phased with SHAPEIT v.2^52^ using default parameters and the HapMap phase II b37 genetic map. Subsequently, the painting profile of each individual was inferred using ChromoPainter. The nuisance parameters n and m were inferred by running ChromoPainter with -in -iM flags for 10 E-M iterations. Given the high computational requirements, the analysis has been carried out only on a subset of the data, using an approach similar to Montinaro et al. 2015^53^. In detail, five (where available, otherwise using all the possible samples) individuals were randomly selected from each population. The inferred parameters were finally used in ChromoPainter specifying all the available samples as donors and recipients. The coancestry matrices based on the length and number of fragments, were then obtained by combining the different chromosomal outputs by means of ChromoCombine. The obtained chunkcount coancestry matrix was harnessed to identify homogeneous groups of individuals using fineSTRUCTURE. In detail, two different runs of 4,000,000 iterations were performed, discarding the first 1 million as burn-in and using a thin interval of 10,000. For each of the clustering approaches, a hierarchical tree was inferred by taking advantage of the “Tree” method and the “maximum concordance state” approach, performing 1 million iterations. In order to assess the robustness of the clustering process, the pairwise coincidence statistics among individuals was evaluated (Supplementary Fig. S6).

### PCA

Principal Component Analysis (PCA) was carried out using coancestry matrix of coping vectors created with ChromoPainter. On the whole, 624 samples were used, including all samples from European, Caucasus and Anatolia and the sub-set of Near Easterners, that on fineSTRUCTURE tree (Supplementary Fig. S4) formed a sister-clade of the European cluster.

### GLOBETROTTER

The time of admixture and mixture profile were estimated for all the previously inferred clusters (targets) using GLOBETROTTER^36^. In detail, the painting profile obtained by ChromoPainter was harnessed by testing for any evidence of admixture using the options null.ind=1 prop.ind=0, and performing 100 bootstrap iterations. For each of the inferred admixture events, only those characterised by bootstrap values for time of admixture between 1 and 400 were considered. Subsequently, the time of admixture was estimated by repeating the same steps with options null.ind=0 and prop.ind=1. For the “non-local” analysis the same procedure has been repeated excluding clusters from the Southern European group as possible sources of the target.

### Projecting ancient Europeans into modern Eurasian variation

In order to contextualise the genetic variation of modern individuals in a European pre-Iron Age context, a PCA analysis was performed in which the ancient genotype data were projected into the PCA inferred on modern Eurasians (Fig. 5A). For ancient samples, genotype data released by Lazaridis et al. 2017^54^ and Olalde et al. 2018^55^ were used, from which non relevant individuals and all the samples with more than the 70% of missingness were removed. After the final filtering 222 samples were retained (Supplementary Table S1B). For modern samples, 568 West Eurasian Individuals were retained. In order to prevent the shrinkage bias in ancient individuals, the “autoshrink: YES” option was used.

### qpADM

The admixture profile of all the analysed populations was reconstructed with qpADM^39^, which harnesses a combination of f4 describing the relationship of “target” and “sources” with a set of outgroups. In detail, the reconstructed admixture profile of each target used any possible four-member combination drawn from a list of putative sources. As a preliminary step, qpWave^56^ was used if: a) the tested sources were significantly different and b) the target may be reconstructed using a specific combination of sources. A p-value threshold of 0.01 was used, as well as the following set of sources (Supplementary Table S1B):

*Anatolia_BA, Anatolia_N, Steppe_EMBA, WHG, EHG, Iran_N, Minoan_Lasithi, Mycenaean, Yorubas, Levant_N, CHG, Europe_EN, Europe_LNBA, Europe_MNChL*

and the following set of Outgroups:

*AfontovaGora3, EHG, ElMiron, GoyetQ116-1, Iran_N, Kostenki14, Levant_N, MA1, Mota, Natufian, Ust_Ishim, Vestonice16, CHG.*

All the tests for which the fitted model was supported were shown in Supplementary Table S5.

## Data Availability

The data for 49 sequences generated in the current study are available in the NCBI-GEO repository through accession nr. GSE129663 and on the Estonian Biocenter website ebc.ee/free_data.

## Acknowledgements

We thank all volunteers who donated their DNA samples. We would like to thank Bayazit Yunusbayev for helpful discussions, Viljo Soo for his help in genotyping and Tuuli Reisberg for assistance in data management. Computational analyses were performed at the High Performance Computing Center of University of Tartu. This research was supported by institutional research funding IUT (IUT24-1) of the Estonian Ministry of Education and Research (ET, EP); the Estonian Research Council grants PUT (PRG243) (EP, MM) and PUT (PUT1339) (AK); the European Union through the European Regional Development Funds with projects No. 2014-2020.4.01.16-0030 (MM, FM) and No. 2014-2020.4.01.15-0012 (MM); the European Union through Horizon 2020 grant no. 810645 (MM); the University of Pavia strategic theme “Towards a governance model for international migration: an interdisciplinary and diachronic perspective” (MIGRAT-IN-G) (OS); the Italian Ministry of Education, University and Research (MIUR): Dipartimenti di Eccellenza Program (2018–2022), Dept. of Biology and Biotechnology “L. Spallanzani”, University of Pavia (AR and OS).

## Author Contributions

M.M. conceived the study. M.M., F.M. and E.T. designed the research. O.S., A.R., J.C., J.D.C., S.M., L.P. and M.M. contributed to sample collection. E.T. and F.M. performed the analyses. E.T., F.M., A.K., M.M. and E.P. interpreted the results. E.T. and F.M. wrote the manuscript with input from all the coauthors.

## Additional Information

The authors declare no competing interests.

