## Supplementary Material for "Genome-wide analysis of Corsican population reveals a close affinity with Northern and Central Italy"

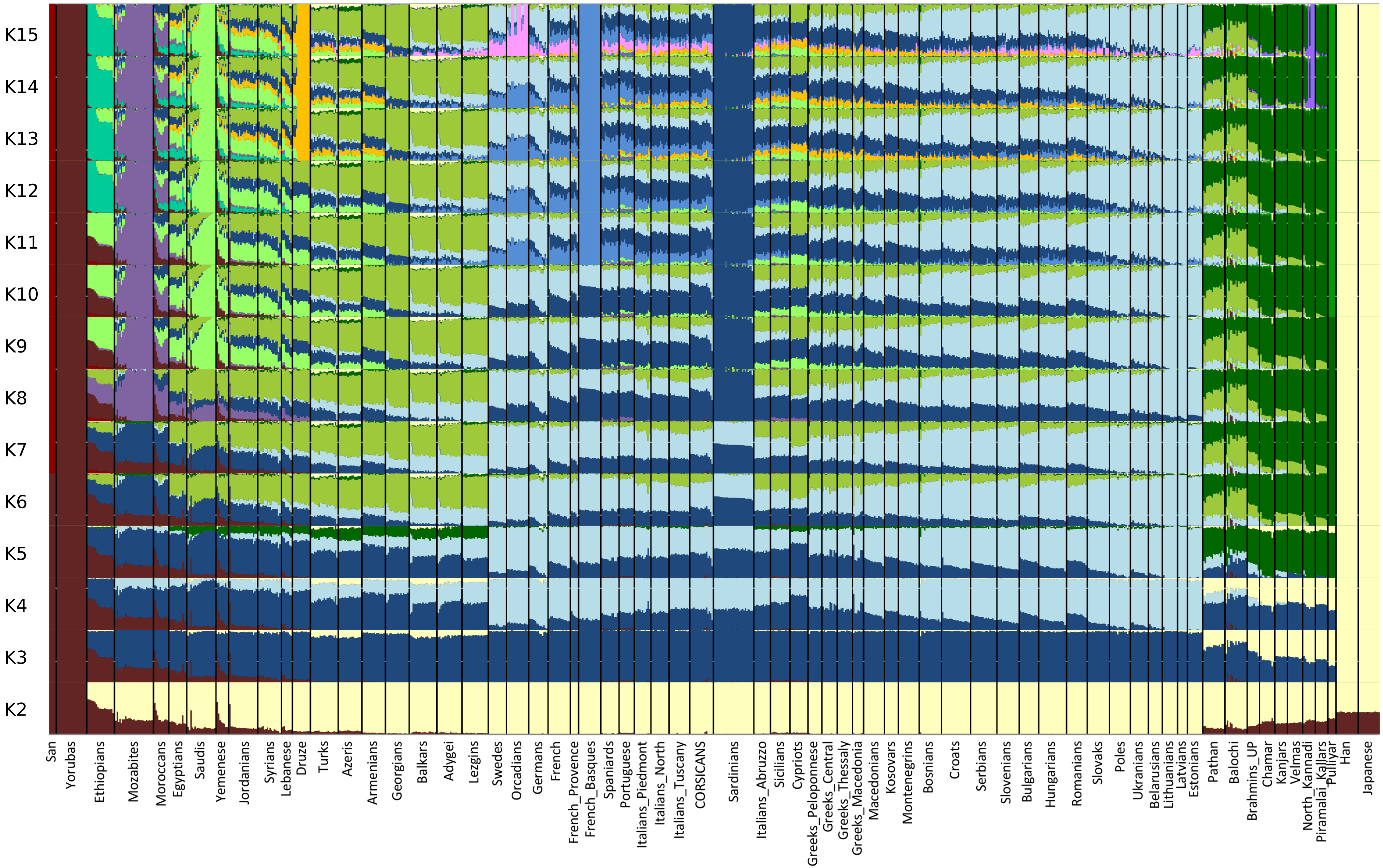


**Supplementary Figure S1.** ADMIXTURE plot of Corsican population in a worldwide context at K =2 to K = 15.

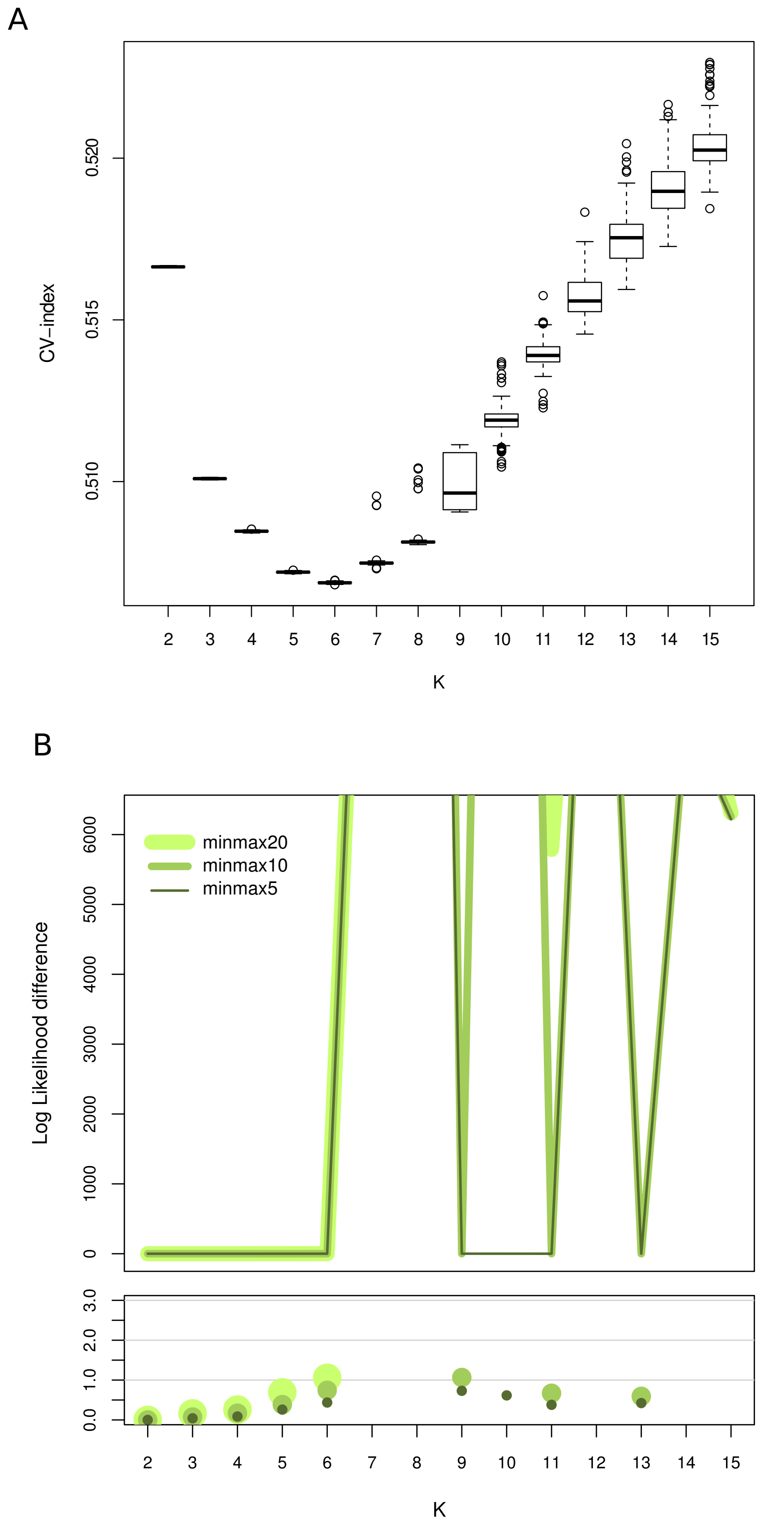


**Supplementary Figure S2.** Selection criteria for optimum level of K.
A) Box and whiskers plot of the cross validation (CV) indexes of all runs of the ADMIXTURE analysis. B) Variation in log-likelihood (LL) scores in the fractions (5%, 10% and 20%) of runs that reached the highest LL values. We assume that at a given K a global LL maximum was reached if 10% of the runs with the highest LL score showed small variation in LL scores.

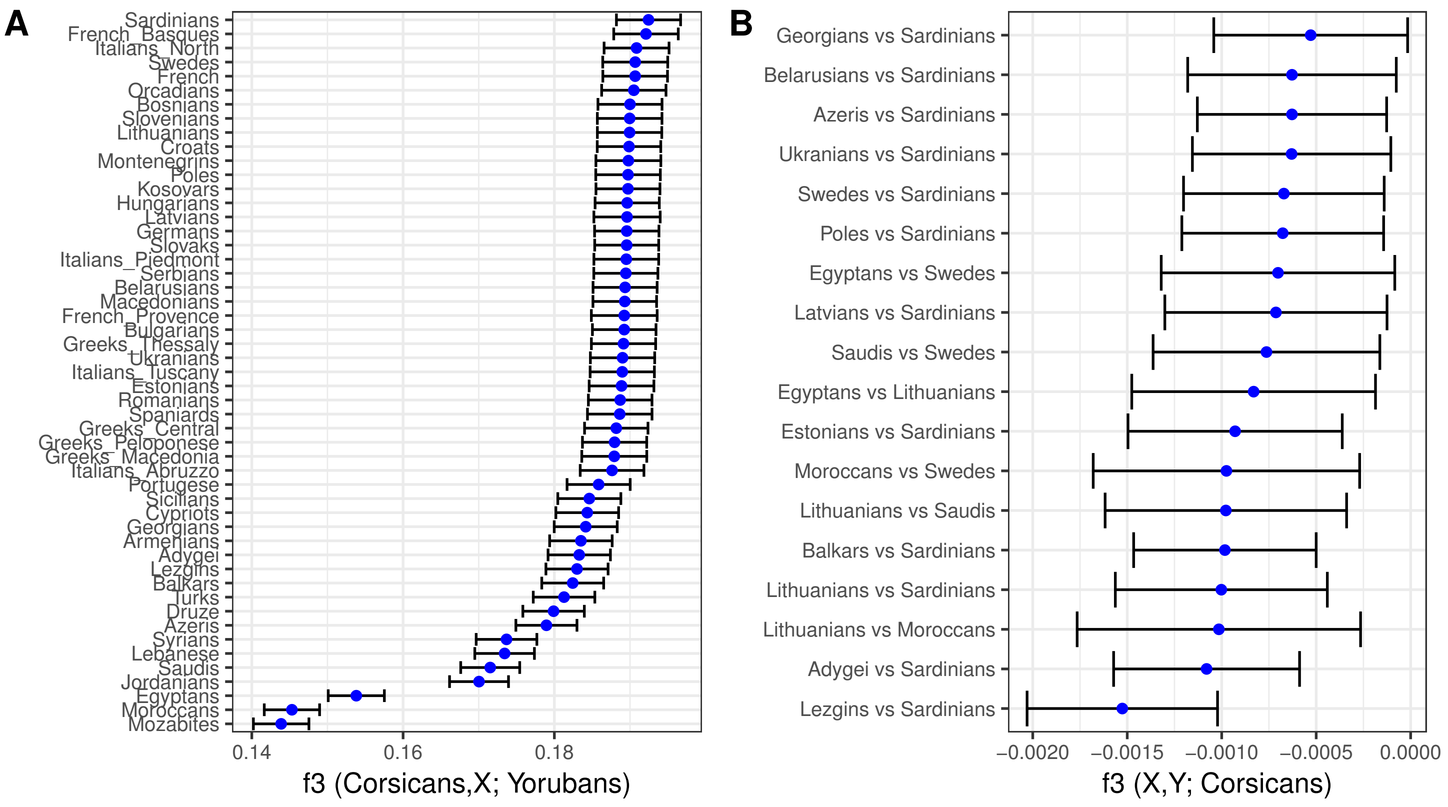


**Supplementary Figure S3.** Results of outgroup f3 and standard f3 tests.
A) Outgroup f3 results of the form f3(Corsicans, X; Yorubans) showing shared drift of Corsicans and reference populations (Supplementary Table S1) from Yoruba populations as outgroup.B) Admixture f3-statistics of the form f3(X ,Y; Corsicans), where X and Y represent all possible pairs of combinations of source populations specified in Supplementary Table S1. Only statistically significant negative results are shown (z-score < -3).

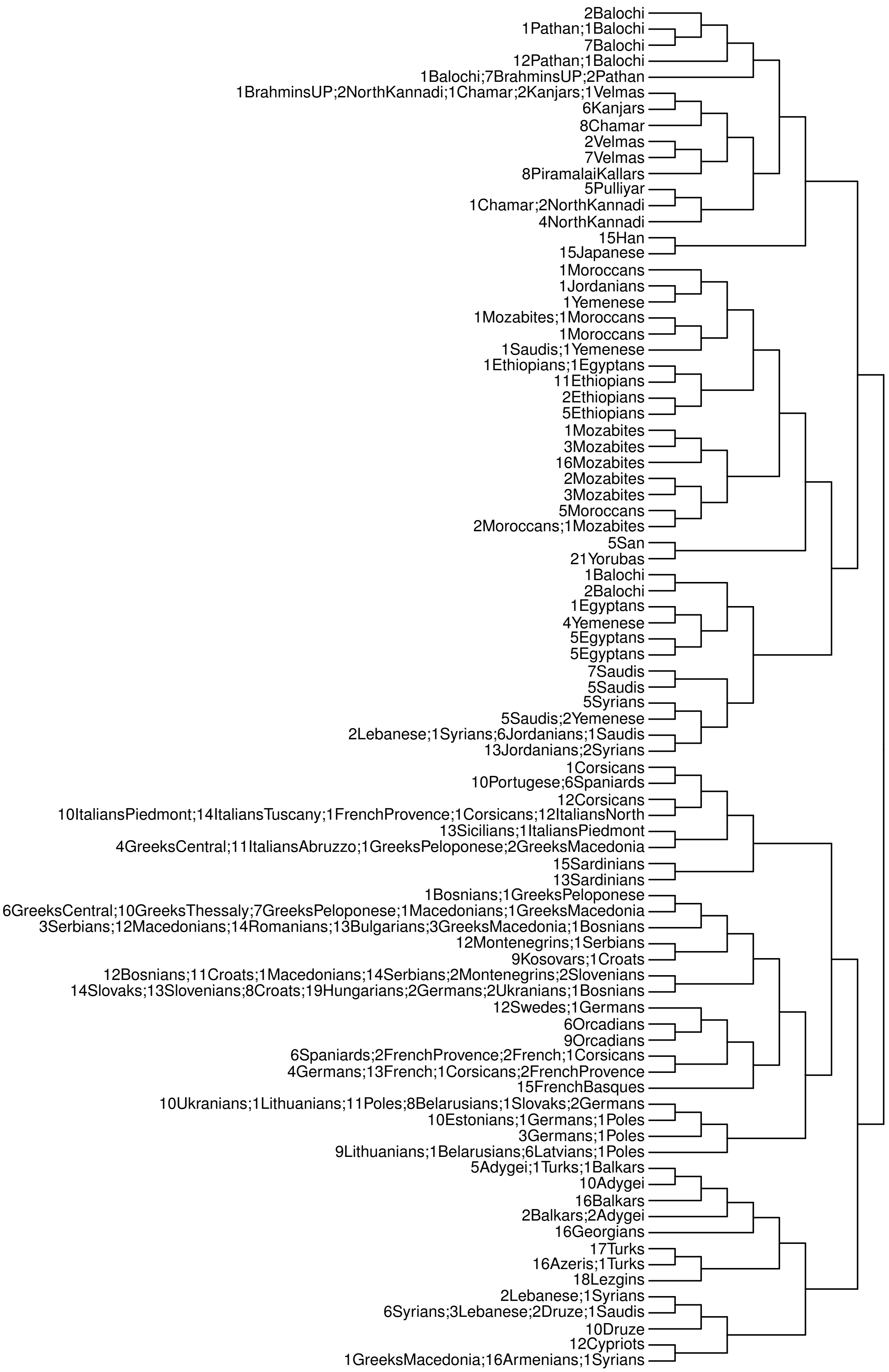


**Supplementary Figure S4.** fineSTRUCTURE dendrogram of all samples. Dendrogram clusters individuals based on similarity of copying vectors. Cluster labels refer to population name and number of individuals from this population. Correspondence with a given cluster name and a macroregional grouping is reported in Supplementary Table S3.

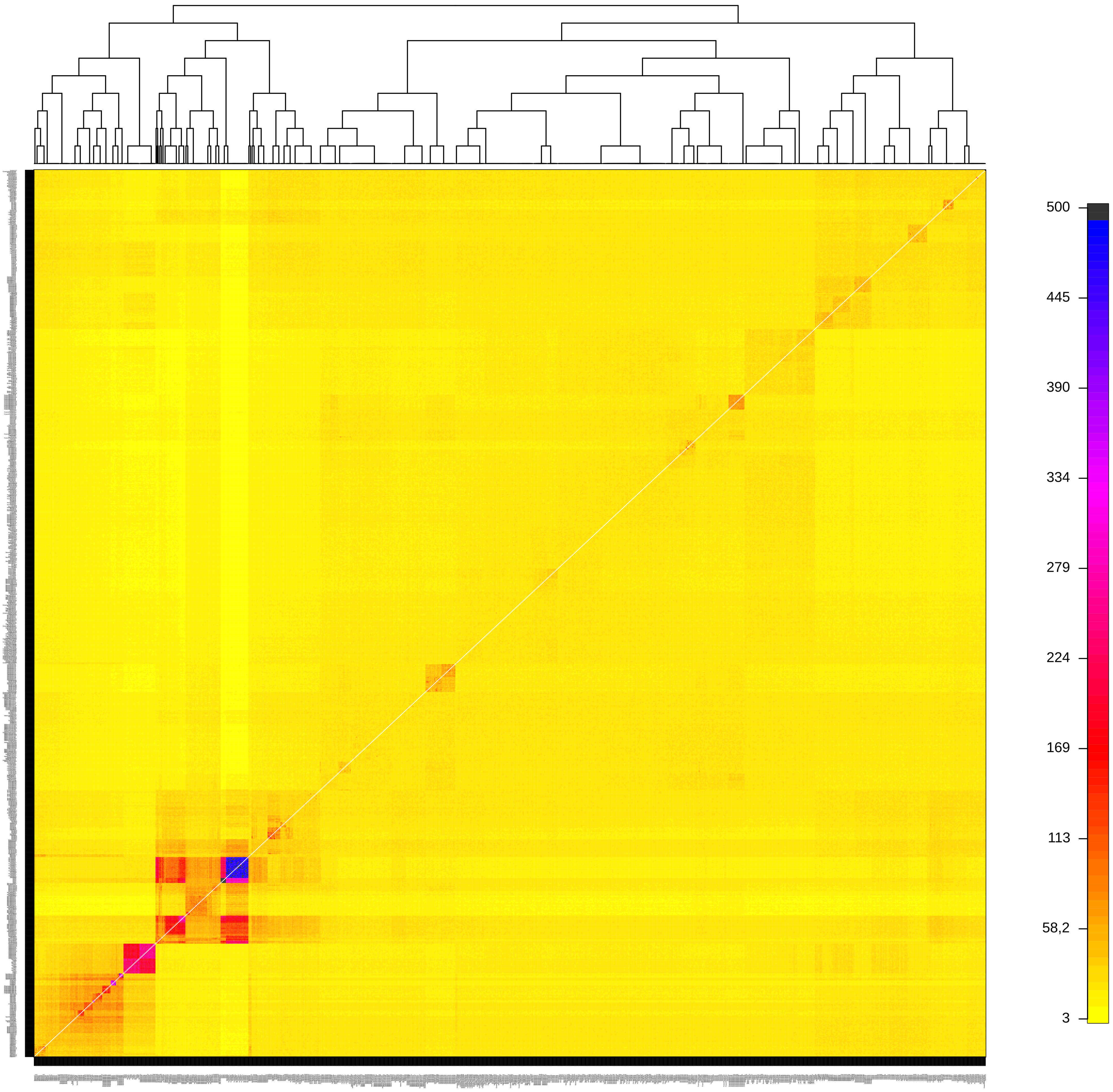


**Supplementary Figure S5.** Coancestry matrix as inferred by ChromoPainter/fineSTRUCTURE.

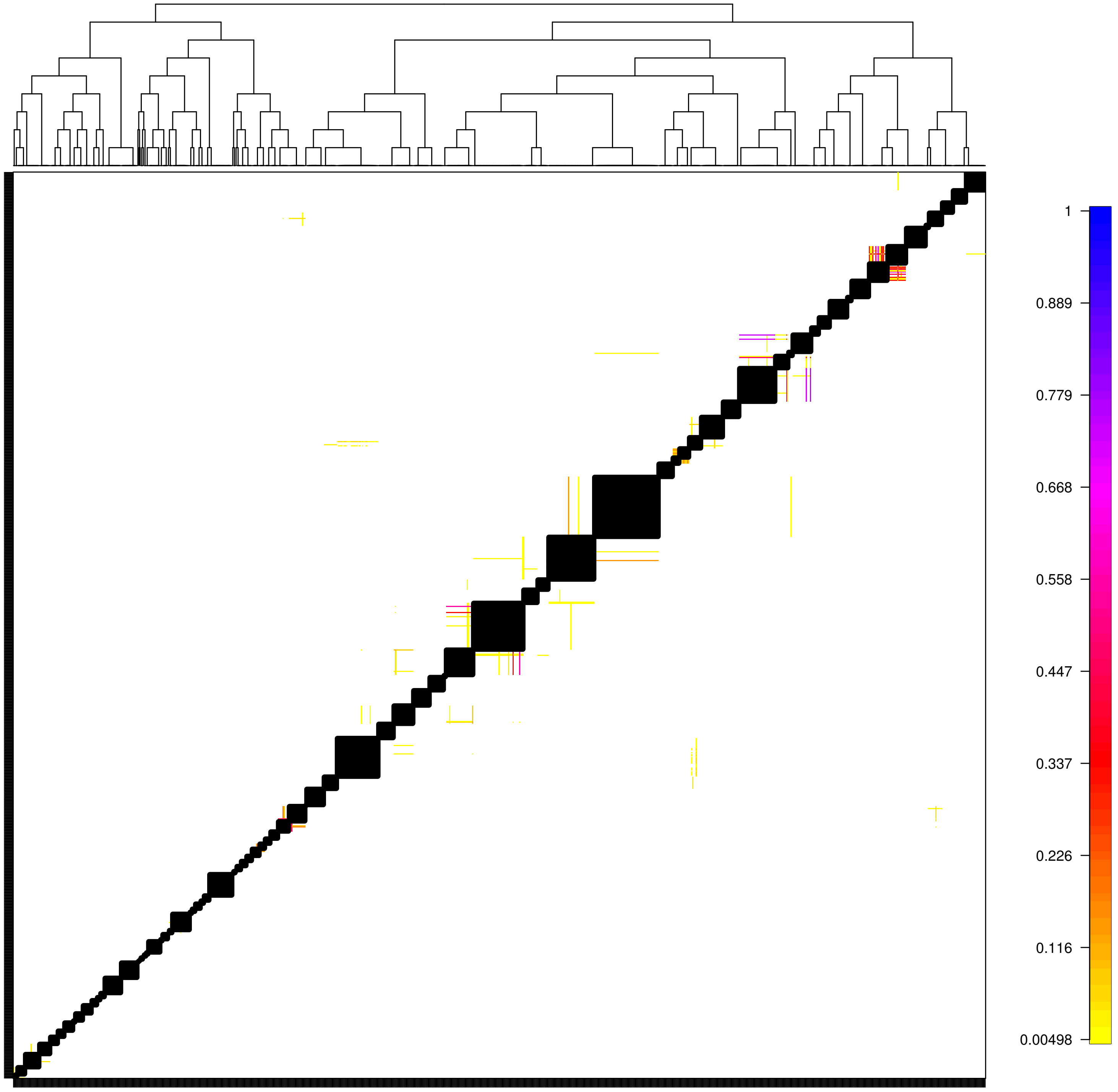


**Supplementary Figure S6.** Pairwise coincidence matrix for the ChromoPainter clustering iterations. The heatmap shows the proportion of ChromoPainter of MCMC iterations for which a pair of individuals fall into the same cluster. As expected, very high values are found at the diagonal, confirming the overall robustness of the clustering approach.

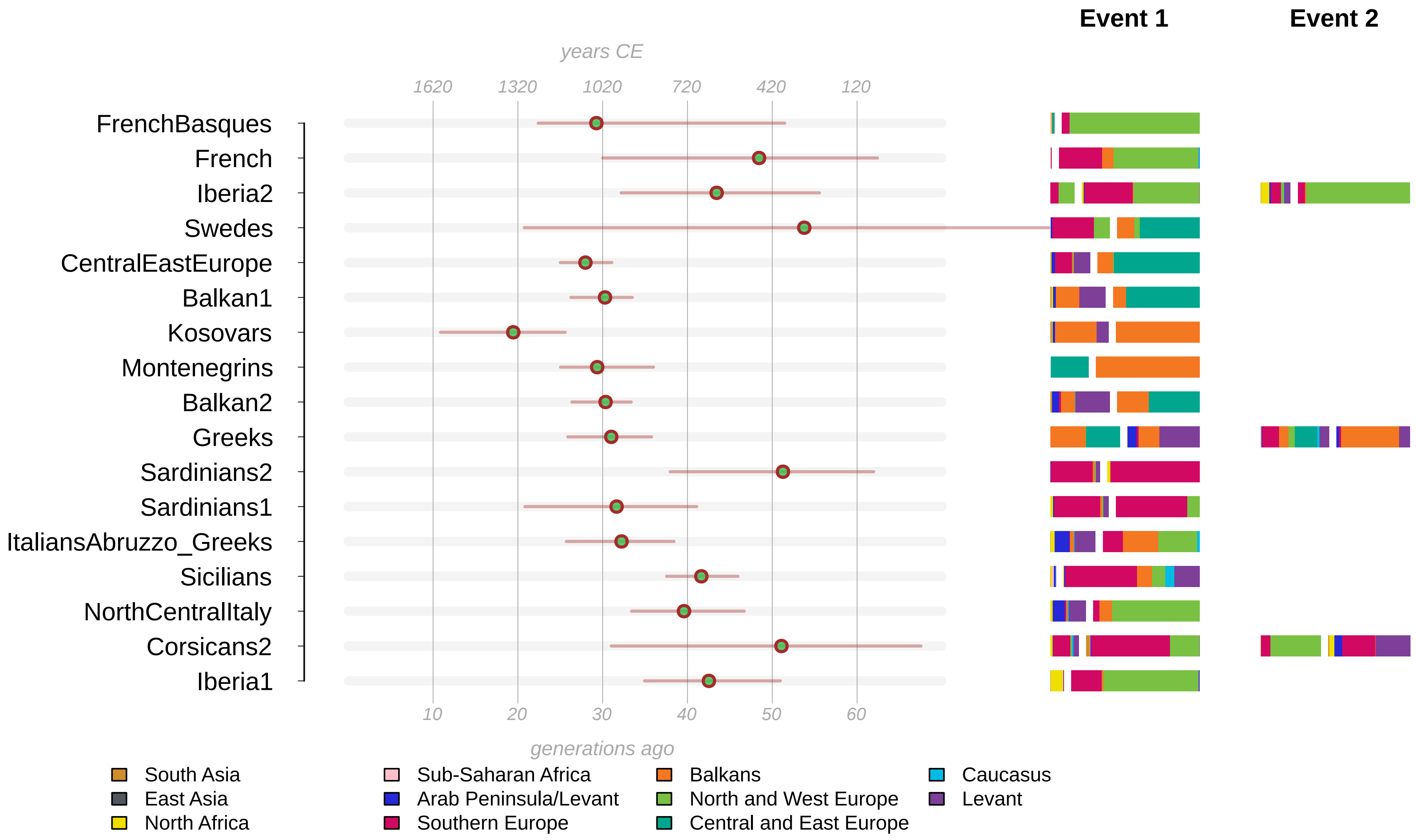


**Supplementary Figure S7.** Admixture dates as inferred by GLOBETROTTER in the “full” analysis. We fit the painting profile of Western Eurasian populations into expected curves for different admixture models, as implemented in GLOBETROTTER. The estimated dates and sources composition are shown.

**Supplementary Tables**

**Table S1.** Overview of dataset composition for different performed analysis. A) Modern populations. B) aDNA samples.

**Table S2.** Pairwise $F_{ST}$ distance among analysed populations

**Table S3.** Inferred clusters by the ChromoPainter/fineSTRUCTURE analysis.

Cluster composition column indicates population name preceded by the numerosity of that population in the cluster.

**Table S4.** Summary of the GLOBETROTTER results for “full” (light blue) and “non-local” (light green) analysis.

**Table S5.** qpAdm results. List of supported four-population scenario for all the analysed populations.
